# Measuring concentration of nanoparticles in polydisperse mixtures using interferometric nanoparticle tracking analysis (iNTA)

**DOI:** 10.1101/2024.01.09.574819

**Authors:** Anna D. Kashkanova, David Albrecht, Michelle Küppers, Martin Blessing, Vahid Sandoghdar

## Abstract

Quantitative measurements of nanoparticle concentration in liquid suspensions are in high demand, for example, in the medical and food industries. Conventional methods remain unsatisfactory, especially for polydisperse samples with overlapping size ranges. Recently, we introduced interferometric nanoparticle tracking analysis (iNTA) as a new method for high-precision measurement of nanoparticle size and refractive index. Here, we show that by counting the number of trajectories that cross the focal plane, iNTA can measure concentrations of subpopulations in a polydisperse mixture in a quantitative manner and without the need for a calibration sample. We evaluate our method on both monodisperse samples and mixtures of known concentrations. Furthermore, we assess the concentration of SARS-CoV-2 in supernatant samples obtained from infected cells.

## Introduction

Measurements of nanoparticle concentrations are important in several fields because they report on the dosage and purity of a given sample. Such information is crucial in drug characterization and administration,^1^ potential toxicity in the food industry,^2^ or in environmental research, where concentrations of unwanted entities such as nanoplgastics in water need to be monitored.^3^ Techniques such as electron microscopy (EM) and atomic force microscopy (AFM) are routinely used for characterization of nanoparticles. While these methods offer excellent size determination, they can generally only yield relative and not absolute concentration values unless the volume of the liquid is controlled to a high degree.^4^ Furthermore, they necessitate a stringent sample preparation procedure based on the deposition of particles on the surface, thus, introducing uncertainties associated with surface wettability and affinity of the particles under study.^5^

Other widely-used techniques for measuring nanoparticle concentration include dynamic light scattering (DLS),^6–8^ nanoparticle tracking analysis (NTA),^6–8^ tunable resistive pulse sensing (TRPS),^7–9^ as well as nanoparticle flow cytometry (nFCM).^8,10^ DLS assesses particle size by analyzing temporal correlations in the light scattered by diffusing particles, whereby the concentration should be low enough to avoid multiple scatterings. ^11^ Generally, nanoparticle concentrations between 10^8^ and 10^12^ particles/ml can be measured in DLS^6^ although the exact range depends on particle size and material.^12^ Since the concentration is extracted from the total amount of the scattered light, reference samples are needed for accurate concentration measurements. Multi-angle DLS (MADLS) does not require reference samples, but the particle refractive index (RI) needs to be known. In NTA, particles in the field of view (FOV) are counted, and this number is converted to a concentration value by using a pre-determined factor. The measured values range between 10^7^ and 10^9^ particles/ml.^6^ Here, the lower limit can be extended by increasing the measurement time, but the upper limit is set by the necessity to avoid intersecting trajectories. In TRPS, particle concentration and size are extracted by counting the particles and estimating the particle volume from the drop of the electric current as it traverses a pore of tunable size. ^9^ For combined size and concentration measurements, a calibration with a sample of known size and concentration is necessary. Previous measurements covered concentrations of 10^8^ to 10^11^ particles/ml ^8,13^ although this range can be adjusted by changing the pressure and the pore size. In nFCM, the light scattered by individual particles is measured as they are sent through a laser beam. A calibration sample is used to relate the scattered light signal to the particle size and the number of particles to concentration.^8^ The concentration needs to be low enough to avoid swarming, where multiple nanoparticles enter the beam simultaneously, but it has to be high enough to prevent the background counts from dominating the measurement.^10^ For a comparison of TRPS, NTA and nFCM, see also Ref. 14.

None of the hitherto available techniques provide an accurate concentration estimate for populations in polydisperse mixtures, especially when the refractive index is unknown or the size ranges overlap.^7^ In a recent work,^15^ we introduced interferometric nanoparticle tracking analysis (iNTA), which performs NTA using interferometric scattering (iSCAT) microscopy.^16,17^ The method features superior performance for determining size and refractive index of nanoparticles and is able to resolve different sub-populations. Previously, we showed that the relative concentrations of subpopulations can be accurately estimated. ^18^ In this work, we present a strategy for performing absolute concentration measurements without the need for a calibration sample. We discuss the theoretical and practical limits of concentration measurements in iNTA and apply the method to supernatants of infected cells, where we specify the concentration change of SARS-CoV-2 virions over time.

### Measurement strategy and experimental procedure

A straightforward approach to the measurement of particle concentrations is to count particles within the FOV of a given volume, depicted schematically as a box in Fig. 1a.^6^ In practice, however, the volume that contributes to the optical signal is not as clearly defined. For example, in case of a FOV defined by a focused Gaussian beam, the extent of the boundaries is not sharp. As a result, whether a particle is counted or not depends strongly on its position and scattering cross section as well as the sensitivity of the setup. This is illustrated by the highlighted section in Fig. 1b. These subtleties call for a particularly careful calibration of the setup on well-characterized samples.

**Figure 1:**
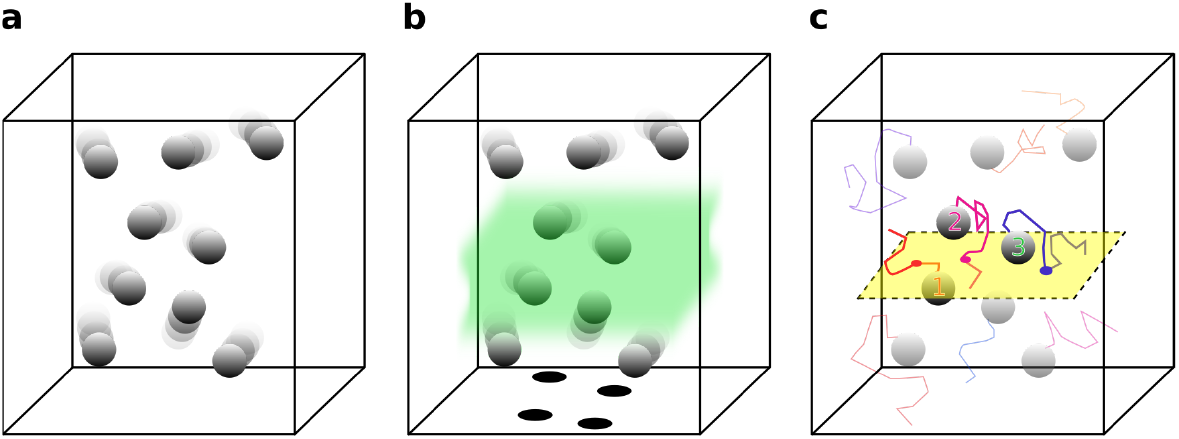
**(a)** A volume of liquid containing a certain concentration of nanoparticles. **(b)** The highlighted region indicates an effective volume that contributes to the optical signal. **(c)** The yellow plane depicts the focal plane. When particles cross the focal plane, their contrast reverses.

In NTA, which uses a dark-field microscope arrangement, the extent of the detection volume in *z* is poorly defined and varies by a large degree with the particle scattering cross section, *i*.*e*., a larger particle remains visible further away from the focal plane than a smaller particle. This results in overestimation of the concentration for larger particles.^8^ To get around this problem, we propose to deduce the concentration by counting the *trajectories* of particles that cross the focal plane. As we show below, this number depends only weakly on the *z*-extent of the detection volume (see Fig. 1c). Implementing this strategy in NTA is nontrivial because the slow change of the PSF along the axial direction makes it difficult to determine the focal plane of a dark-field microscope with great precision. In iSCAT microscopy, the central lobe of the PSF approaches the diffraction limit and the signal undergoes a contrast inversion between maximally bright and maximally dark when the particle crosses the focal plane.^19^ This feature allows us to perform a robust measurement of the nanoparticle concentration.

The optical setup was described in a previous publication.^15^ In the current measurements we used a different microscope objective (Leica HC PL APO 160x 1.43 Oil), yielding a pixel size of 71 nm and a FOV of 5.2 *×* 5.2 μm^2^, whereby the microscope focus was set at about 1 μm above the coverglass. A uniform illumination was achieved by employing acousto-optical deflectors (AOD) in the incident beam path, scanning about 10x faster than the acquisition rate. Measurements were performed at 10 kHz with 50 μs exposure time. The measurement and analysis procedures are the same as described previously.^15^ We recorded two sets of 300 one-second-long videos of particles diffusing in 100 μL volumes of fluid inside individual ibidi wells using pylablib cam-control.^20^ Recorded videos were analyzed by applying median background correction and radial variance transform.^21^ Particles were tracked using the trackpy python package^22^ with linking radius of 8 px = 565 nm. Particles were allowed to disappear for at most 20 frames before the trajectory would get a new identifier. Only particles whose central lobe at the position of the maximum contrast could be fitted with a Gaussian function with standard deviation between 100 and 120 nm were considered for further analysis, as we have found this to be the range in which particles crossing the focal plane in monodisperse samples reach their maximum contrasts.

## Simulations

In order to relate the number of detected trajectories to an absolute particle concentration, we simulated a single particle with a given diffusion constant in a 10*×*10*×*10 μm^3^ box, corresponding to a particle concentration of 10^9^ particles/ml. Particle diffusion was simulated for 0.1-100 s with a random starting position at a frame rate of 10 kHz. Simulations were repeated 30,000 times. The average number of trajectories longer than a given threshold per video was extracted to be related to particle concentration. Figure 2a displays the *x* − *z* projections of five exemplary trajectories of diffusing particles with *D* = 6 μm^2^/s over 1 s. The dashed gray line indicates the position of the focal plane. In Fig. 2b, we crop the trajectories in (a) to the detection volume (here -3 μm *< x, y <* 3 μm and *z <* 3 μm), indicated in gray. As in our data analysis, if the particle leaves the detection volume for more than 20 frames, it is assigned a new identifier and counted as a new trajectory. We only keep the particles that cross the focal plane. This results in seven trajectories, each marked by a different color in Fig. 2c, which we would analyze in our experiment. As in the analysis of the experimental data, we impose a condition that only trajectories with more than 100 localizations are considered.

**Figure 2:**
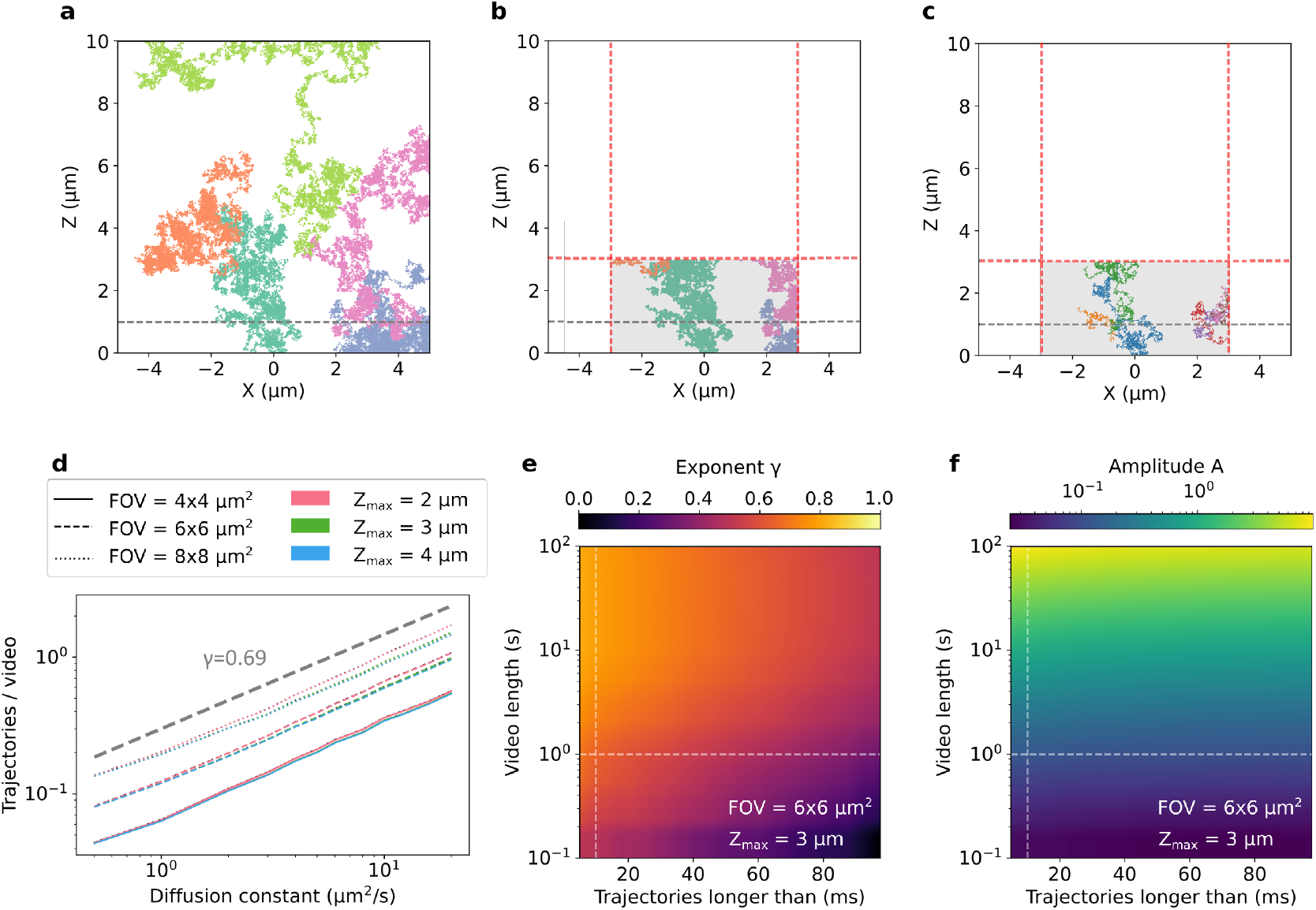
**(a)** Projections of trajectories from five exemplary particles with diffusion constant of 6 μm^2^/s diffusing in a 10 *×*10 *×*10 μm^3^ box over 1 s. Gray dashed line shows the position of the focal plane. **(b)** Same as (a), but only the parts of the trajectories in the detection region (shaded in gray: 6 *×* 6 μm^2^ FOV and *z <* 3μm) are shown. Red lines indicate the boundaries. **(c)** As a particle leaves and enters the detection value, it receives a new identifier. The resulting seven trajectories that would be detected in an experiment are shown with different colors. **(d)** Average number of trajectories longer than 10 ms (100 localizations) detected in 1 s-long video is plotted vs. diffusion constant for different FOVs and *z*_max_. The data can be fitted with the power law of the form *T* = *AD*^*γ*^. Thick gray line shows *T ∝ D*^0.69^. **(e)** The extracted value of power law exponent *γ* for different video lengths and different minimal trajectory lengths. Here, we assume 6 *×* 6 μm^2^ FOV with *z*_max_=3 μm. The dashed lines indicate the parameters used in the experiment: 1 second long videos with trajectories longer than 10 ms. **(f)** Same as (e), but now the power law amplitude (*A*) is plotted.

Figure 2d shows the average number of detected trajectories with more than 100 localizations recorded in a 1 s long video. Here, we assume 10^9^ particles/ml and consider different sizes of FOVs and values of *z*_max_. We note that the average number of the detected trajectories depends strongly on the diffusion constant of the particles as well as on the FOV size. The dependence on *z*_max_ is very weak because only particles that cross the focal plane placed 1 μm above the coverglass are considered. We find that the average number of trajectories follows the power law *T* = *AD*^*γ*^, where *T* is the number of trajectories, *D* is the diffusion constant, *A* is the proportionality constant, and *γ* is the exponent. The value of the exponent is independent of the FOV size and *z*_max_. We then explored the dependence of the fitting parameters *γ* and *A* on the video length and minimal trajectory length. In Fig. 2e,f, we see that *γ* varies strongly with both, while *A* is predominantly determined by the video length. The frame rate affects the results as well. Therefore, it is important to conduct simulations with the experimental parameters corresponding to the setup. For our parameters (5.2 *×* 5.2 μm^2^ FOV, 10 kHz imaging rate, minimum trajectory length of 100 points), we obtain *A* = 0.1 and *γ* = 0.69, which we will use in the following sections.

## Results and discussion

### Benchmarking with monodisperse samples of known concentration

To benchmark our methodology, we studied different dilutions of monodisperse particles that were characterized by the manufacturer. Particles with a density much larger than water (*e*.*g*., gold or silica nanoparticles) may bias concentration measurements due to sedimentation. To avoid this systematic issue in our benchmarking measurements, we chose polystyrene (density of 1.05 g/cm^3^). We measured 13 dilutions of NIST-certified 40 nm, 60 nm and 100 nm polystyrene spheres (PS) for five minutes twice (300 videos each time). In each case, we extracted the number of detected trajectories that contained more than 100 localizations. Figure 3a plots the outcome versus the expected particle concentration. We note that the number of trajectories increases with the expected particle concentration (*C*_exp_) until around 10^11^ particles/ml, where the sample becomes too crowded for reliable particle tracking.

**Figure 3:**
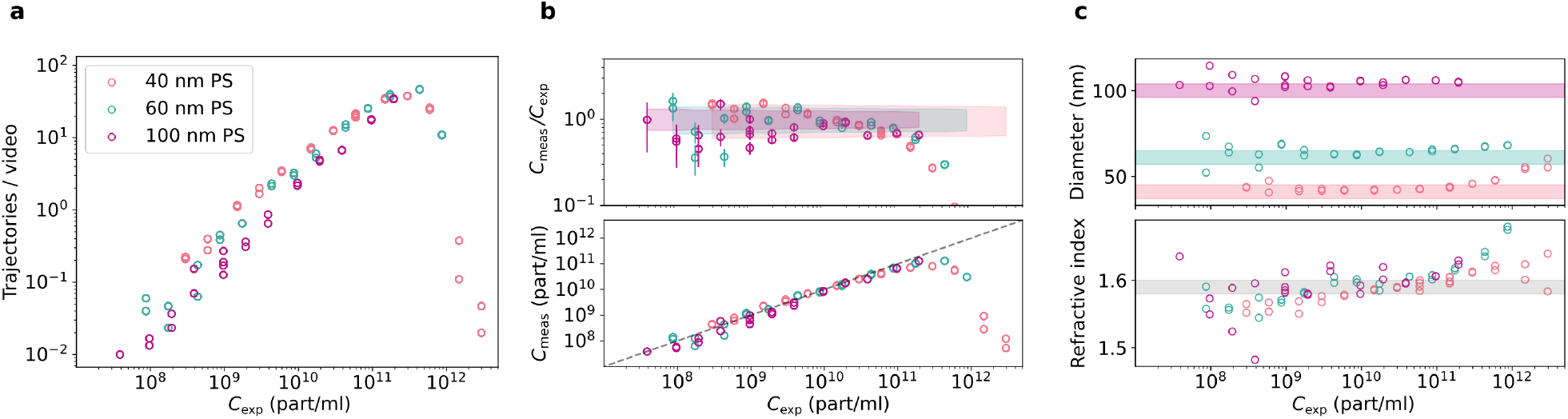
**(a)** Average number of trajectories recorded during 300 one-second long videos for 40 nm (red), 60 nm (green) and 100 nm (violet) polystyrene spheres (PS) for different expected particle concentrations. Trajectories longer than 100 localizations were analyzed. **(b)** Upper panel: ratio of measured to expected concentrations. Shaded region indicates the limits on particle concentration according to the data provided by the manufacturer. Lower panel: Concentration in particles per ml measured using the data in (a). **(c)** The extracted particle size and RI. Shaded regions indicate manufacturers’ specifications.

The lower panel of Fig. 3b shows the measured particle concentration (*C*_meas_), calculate from trajectory number and diffusion constant as described above. The dashed line has a slope of 1. We examine the agreement between the measured and expected concentration values by considering the ratio of the two, as shown in the upper panel. The error bars show the statistical error due to the limited number of trajectories, calculated as 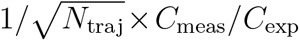. The colored areas indicate the range of possible deviations due to the uncertainty in stock concentration and sample dilution. We find a good agreement between the measured and expected concentrations up to 10^11^ particles/ml. In Fig. 3c, we show the median extracted diameter and the median refractive index as a function of the expected particle concentration. Above 10^11^ particles/ml, the crowdedness of the sample biases the detection towards larger particles with larger RI. For low concentrations, the small number of recorded trajectories lead to an increase in the statistical error in concentration, diameter and RI determination.

### Polydisperse samples with known concentrations

Next, we determined the concentrations of populations in a mixture of particles with overlapping sizes. Here, we used 100 nm PS and silica beads (SB), whereby the latter had a broad size distribution, as we previously reported in Ref. 15. We measured both types of particles with three different dilutions, resulting in nominal concentrations of approximately 10^9^, 10^10^ and 10^11^ particles/ml. For the case of PS sample, we see that the measured concentration agrees well with its nominal value (Fig. 4a bottom). For the SB sample, the nominal concentration of the stock solution was not known. Our measurements provided an estimate of 1.75 *×* 10^13^ particles/ml. Assuming this stock concentration, the measured and nominal concentration values agree well for all three dilutions (Fig. 4b bottom).

**Figure 4:**
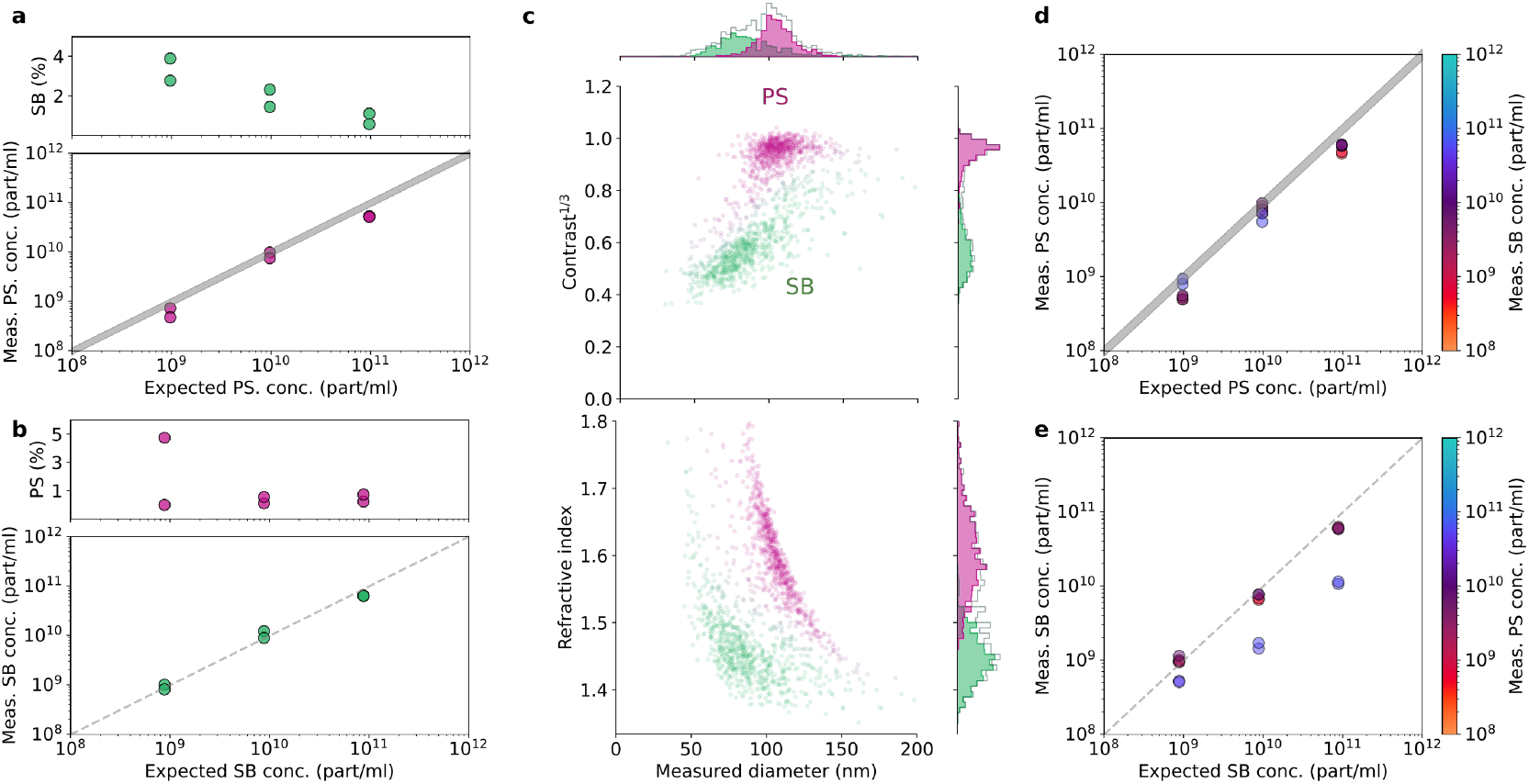
Determining subpopulation concentrations for a mixture of PS and SB. **(a)** Bottom: measured versus expected concentrations for 100 nm PS. Gray area is shaded using the values provided by the manufacturer. Top: percentage of particles misclassified as 100 nm SB when random forest classifier was applied. **(b)** Same as (a) but PS and SB are switched. **(c)** An exemplary 5 minute measurement of a 1000x dilutions of PS and SB mixed in 1:1 ratio. The top plot shows third root of iSCAT contrast plotted vs. particle size, while the bottom plot show the refractive index plotted vs. particle size. **(d)** The measured vs. expected concentration of PS in mixtures of PS and SB. Color indicates the measured concentration of SB. **(e)** Same as (d) but PS and SB are switched.

To examine the robustness of our assignments, we trained a random forest classifier on a subset of the data^23^ and used that classifier to estimate the percentage of the particles that become misclassified in pure samples. We only consider particles for which the confidence of belonging to a group is higher than 90%. This approach eliminates on average about 12% of all particles. We find that the percentage of the misclassified particles decreases with increasing particle concentration and always remains below 5%.

We measured 9 different mixtures of PS and SB (combinations of 1-1 mixtures of 100x, 1000x and 10000x dilutions of both samples) and applied our classifier to the results. The 2D plots of the size and the third root of the iSCAT contrast (top) or RI (bottom) are shown in Fig. 4c for an exemplary sample (1000x diluted PS:1000x diluted SB). We calculated the absolute concentration of particles according to *C*_X_ = *C*_X,90_*/*(*C*_PS,90_ + *C*_SB,90_) *× C*_total_, where X stands for PS or SB, and *C*_X,90_ indicates the concentration of the particles for which classifier confidence was above 90%. The plots of the measured PS (SB) concentrations are displayed in Fig. 4d (e), where in each case the color indicates the measured concentration of the other species. We point out that the presence of SB particles does not seem to affect concentration measurements of the PS components. However, high PS concentrations lower the accuracy in the measurement of SB concentration (blue points in Fig. 4e). This is because the iSCAT contrast of PS is 2-8 times greater than that of SB, so that their presence creates a background against which SBs are difficult to distinguish.

### Virus concentration measurements

Assessment of the concentration of viral particles in medical samples is often a nontrivial task.^24^ To demonstrate the application of our methodology to uncharacterized medical species, we examined samples obtained from cells that were infected with SARS-CoV-2. We collected the supernatant of the infected cells at several time points (see Methods) and found that almost no particles were detected for the first 12 hours post-infection (hpi). In Fig. 5a, we present the results of 10-minute long measurements of 12, 18, 24 and 48 hpi samples. We observe a steady increase in the number of particles over time, which can be clustered into two separate populations. Moreover, we note that the relative ratio of the two populations changes with time. By applying a Gaussian mixture model (GMM) with full covariance, ^23^ we differentiated the two populations quantitatively as plotted in Fig. 5b. We find that after 12 hpi the concentration of the high-contrast population (teal) grows monotonically, while the concentration of the low-contrast population (pink) remains constant. We speculate that the former may be virus particles released by the infected cells. We find that the hydrodynamic diameter of the particles in the high-contrast population is between 90 and 140 nm, which is larger than the reported diameter of the lipid bilayer of SARS-CoV-2 as measured with Cryo-EM (91 *±* 11 nm).^25^ Nevertheless, the measured hydrodynamic diameter is reasonable if we account for the size of the S-protein (length *∼* 22 nm^26^), which is present on virions in various quantities. Assuming that the population with the higher contrast corresponds to SARS-CoV-2 particles, we hypothesize that the second population with a lower contrast may represent extracellular vesicles (EVs) and or protein aggregates released by the cells.

**Figure 5:**
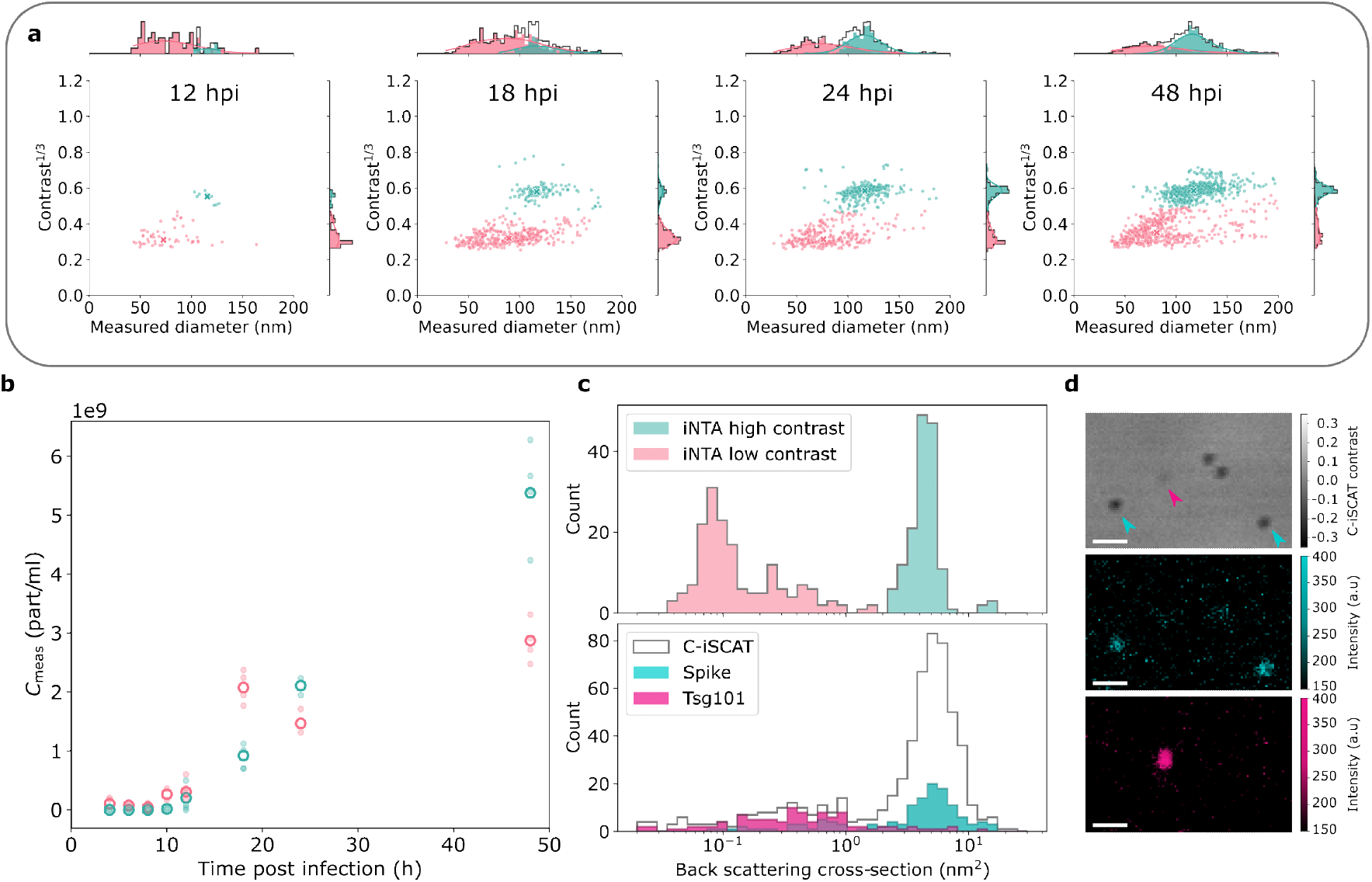
Measurements of SARS-CoV-2 concentration in supernatant of infected cells over time. **(a)** Results of 10 minute iNTA measurements of samples with different time post-infection, as indicated. Teal and pink colors indicate different populations as extracted by the GMM. **(b)** Concentrations of particles as a function of time after infection for the two populations. **(c)** Upper: iSCAT contrast extracted from the iNTA data recorded for 24 hpi sample converted to back-scattering cross-section. Lower: C-iSCAT contrast converted to back-scattering cross-section. Colors indicate presence of fluorescence labels for S-protein (teal) and Tsg101 (magenta). **(d)** Exemplary C-iSCAT and fluorescence images for particles adsorbed on a cover glass. Arrows in the top panel indicate particles positive for S-protein (teal) or Tsg101 (magenta). Scale bar is 1 μm.

To check our assignment of the two observed populations, we performed combined C-iSCAT and fluorescence microscopy, as detailed in the Methods section.^27^ Briefly, we immunofluorescently labeled particles from the supernatants for S-protein (Spike, SARS-CoV-2 marker) and Tsg101 (EV marker) and measured their contrast and fluorescence signal. We then used PS data as reference to convert the C-iSCAT contrast to a scattering cross section which was adjusted to account for the different measurement wavelengths in C-iSCAT and iNTA. The top panel in Fig. 5c shows the particle back-scattering cross section extracted from the iNTA measurements for the 24 hpi sample. The corresponding histogram of back-scattering cross sections obtained using C-iSCAT also reveals two distinct peaks, as shown in the bottom panel of Fig. 5c. By exploiting the simultaneously acquired fluorescence intensity, we can now identify the two populations. The particles with a higher mean back-scattering cross section of *∼* 3 *−* 10 nm^2^ correspond to particles positive for S-protein (teal), while the particles with a smaller mean back-scattering cross section of *∼* 0.1 *−* 5 nm^2^ are predominantly Tsg101-positive and are, therefore, classified as EVs (magenta). An representative FOV is shown in Fig. 5d, revealing the presence of S-protein positive particles (middle panel) as well as Tsg101 positive particles (bottom panel). The combination of C-iSCAT and fluorescence measurements confirms that the high-contrast population can, indeed, be considered as SARS-CoV-2 particles produced by the infected cells.

In the top panel of Fig. 5d, we note additional unlabeled particles with high contrast. We attribute these to virions without the fragile S-proteins that easily shear off the virus particles or incomplete immunofluorescence labeling. Moreover, we observe a population with scattering cross section below 0.2 nm^2^ (see Fig. 5c), which may correspond to protein aggregates.^15^ The fact that we do not detect these aggregates in C-iSCAT could be because they are less likely to adhere to the surface or their signal is not distinguishable from the cover glass roughness.

## Conclusions

We have demonstrated that iNTA is a useful tool not only for characterization of particle size and refractive index but also for a quantitative assessment of the particle concentration. Moreover, concentrations of different populations in polydisperse samples can be reliably determined, even if sizes overlap, which to our knowledge had not been achieved before. In particular, we presented an approach for measuring concentrations without the need for tedious calibration procedures. We find good performance for concentrations up to 10^11^ particles/ml. The lower limit is about 10^8^ particles/ml for a 5 minute measurement, but can be improved by increasing the measurement time. We showcased the power of our technique by determining the concentration of virus particles and residual extracellular vesicles obtained from cells infected with SARS-CoV-2.

## Methods

### Nanoparticle specifications

The following types of nanoparticles were used:

- 40 nm polystyrene: Thermo Fisher, Cat. No.: 3040(-004), Lot: 230327, Certified Mean Diameter: 41nm ± 4nm, k=2 [PCS], Coefficient of Variation: not determined, RI: 1.59 @589nm; Density: 1.05 g/cm3; Approximate concentration: 1% solids
- 60 nm polystyrene: Thermo Fisher, Cat. No.: 3060(-008), Lot: 228547, Certified Mean Diameter: 61nm ± 4nm, k=2 [TEM], Coefficient of Variation: 15.6%, RI: 1.59 @589nm; Density: 1.05 g/cm3; Approximate concentration: 1% solids
- 100 nm polystyrene: Thermo Fisher, Cat. No.: 3100(-008), Lot: 229003, Certified Mean Diameter: 100nm ± 4nm, k=2 [TEM], Coefficient of Variation: 7.7%, RI: 1.59 @589nm; Density: 1.05 g/cm3; Approximate concentration: 1% solids
- 100 nm silica: Corpuscular, Cat. No.: 140120-10, Lot: NX731, Mean Diameter: 101.9nm, Polydispersity index: 0.02

This information was used to calculate the stock concentration of PS to be 2-3.6 *×*10^14^ particles/ml, 6.6-9.8 *×*10^13^ particles/ml and 1.6-2.1 *×*10^13^ particles/ml for the 40 nm, 60 nm and 100 nm PS, correspondingly. For the silica beads (SB), it was not possible to calculate the stock concentration *a priori*.

### Sample preparation

Chambered cover glasses (ibidi *μ*-Slide 18-well with glass bottom) were used for iNTA measurements. The chambered cover glasses were plasma cleaned (1 minute in oxygen plasma at 500 W). Next, 60 μl of 40 nm GNPs from BBI Solutions diluted at 1:200 ratio in milliQ water were introduced into one of the wells. The chambered coverglass was then placed on a heating plate until the liquid evaporated. The remaining 40 nm GNPs on the surface served as a reference to set the focus correctly. Samples were kept covered prior to measurement in order to avoid contamination.

For measurements on SARS-CoV-2, the chambered cover glasses were additionally passivated. In order to do that, the coverg lasses were first plasma cleaned (1 minute in oxygen plasma at 500 W). Wells were filled with 100 μl of 10 mg/mL mPEG2000-Silane dissolved in PEG solution (95% Ethanol (v/v), 5% milliQ, pH was set to 2.0 with 1 M HCl). The chambered coverglass was then incubated at 50 °C. Once the solution fully evaporated, the chambered coverglass was sonicated for 10 minutes in milliQ water and blow dried with nitrogen gas. Passivated coverg lasses were used the same day.

For monodisperse samples, NIST-certified polystyrene spheres (PS) of diameter 40 nm, 60 nm and 100 nm were diluted in milliQ water with dilutions between 100x and 10^6^x. The resulting samples were stored at 4 °C until the measurement. For mixtures, 100x, 1000x and 10000x dilutions of PS and silica beads (SB) were mixed in a ratio of 1:1 to form a total of 9 mixtures.

SARS-CoV-2 samples were prepared from supernatants of infected Vero E6 cells at different time points. Vero E6 cells were grown in 6-well plates to confluency and infected at an MOI of 3 with SARS-CoV-2, an isolate from 2020 kindly provided by the University Hospital Erlangen. Cells were incubated with the virus for 20 min, washed once with phosphate-buffered saline (PBS) and then incubated with 2 ml optipro medium (Thermo Fisher) with added glutamine (Thermo Fisher). After 0, 4, 6, 8, 10, 12, 18, 24, and 48 hours, the supernatants were collected, centrifuged at 3000 g for 30 minutes to remove cells and stored at -80 °C. For iNTA measurements, samples were thawed and inactivated for 2 hours at room temperature by adding PFA (EMS) to a final concentration of 4 % (v/v). Inactivated samples were stored at 4C until measured. For immunofluorescence measurements, 100 μl of inactivated SARS-CoV-2 samples were incubated 90 min at room temperature in glass-bottom dishes (ibidi), unbound particles removed, samples blocked with 4 % BSA (w/v) in PBS and incubated over night at 4 °C with SARS-CoV-2 Spike Protein S2 mouse monoclonal antibody 1A9 (Thermo Fisher) and TSG101 rabbit polyclonal antibody 14497-1-AP (Proteintech) diluted 1:1000. Samples were washed 3 times for 5 minutes with PBS and incubated for 2 hours at room temperature with secondary antibodies anti-mouse AF488 and anti-rabbit AF561 (Thermo Fisher) diluted 1:1000. C-iSCAT and fluorescence images were acquired in PBS.

### C-iSCAT measurement setup, measurement and data analysis procedure

The confocal iSCAT (C-iSCAT) setup was recently described.^27^ Briefly, for C-iSCAT measurements, the laser illumination wavelength was 445 nm, and a 20:80 (R:T) beam splitter and a 450/50 nm bandpass filter were used. Confocal fluorescence microscopy was performed by illuminating the sample with a 488 nm laser beam and use of a 525/50 nm bandpass filter in detection as well as a 561 nm laser beam with a 595/50 bandpass filter in detection. The effective voxel size was set to 30 *×* 30 *×* 30 nm^3^, the FOV was approximately 30 *×* 30 μm^2^, and the total z-range was about 900 nm. The pixel dwell time was 4 μs. The measurement and analysis procedures were previously described.^27^ After acquiring an individual z-stacks, a low-pass filter using a Gaussian distribution with a kernel size of 25 pixels (750 nm) was applied to each z-plane to determine the background intensity, *I*_bg_. Thus, the C-iSCAT contrast, *C*_C*−*iSCAT_ = (*I*_det_ *− I*_bg_)*/I*_bg_, was calculated in each z-plane. A radial variance transform^21^ was applied to detect particles in each background corrected z-plane. The minimal radius was set to 1 pixel, and the maximal radius was set to 4 pixels. The localization of the particles was performed using the trackpy python package^22^ with a radius of 7 pixels and a minimum mass of 1.2 for the RVT signal. To extract the C-iSCAT contrast as well as the fluorescence intensity, we calculated the mean of the central three pixels around the central maximum of the obtained particle localizations. The SNR of the fluorescence signal was improved by performing a maximum intensity projection prior to extracting the individual intensity values in each channel. We determined a threshold for the fluorescence intensity detection based on the mean value of the whole FOV and the standard deviation thereof. Thus, we obtained a threshold of 150 counts for both channels. In order to calibrate the C-iSCAT contrast, we measured 40 nm, 60 nm, and 100 nm PS nanoparticles as specified above. We performed the contrast-to-back-scattering cross section calibration analogously to the iNTA calibration.^15^ We set the refractive index for polystyrene at the *λ* = 445 nm illumination wavelength to be *RI*_PS_(445 nm) = 1.6148 and of the medium *RI*_medium_(445nm) = 1.3372. From this, we derived the calibration factor *β*_C*−*iSCAT_ = 3.0 *×* 10^7^ m^*−*1^.

## Acknowledgement

The authors thank S. Ilhoff for technical assistance, as well as S. Jiang, H. Lee, M. Mazaheri and R. Taylor for careful reading of the manuscript and insightful comments. We also acknowledge support from the Institute of Virology at the University Hospital of Friedrich-Alexander University in Erlangen. A.D.K. also thanks Christiane Nüsslein-Volhard-Stiftung Fellowship. This work was financed by the Max Planck Society. The left panel of the TOC Graphic was created using DALL-E 3.

## TOC Graphic

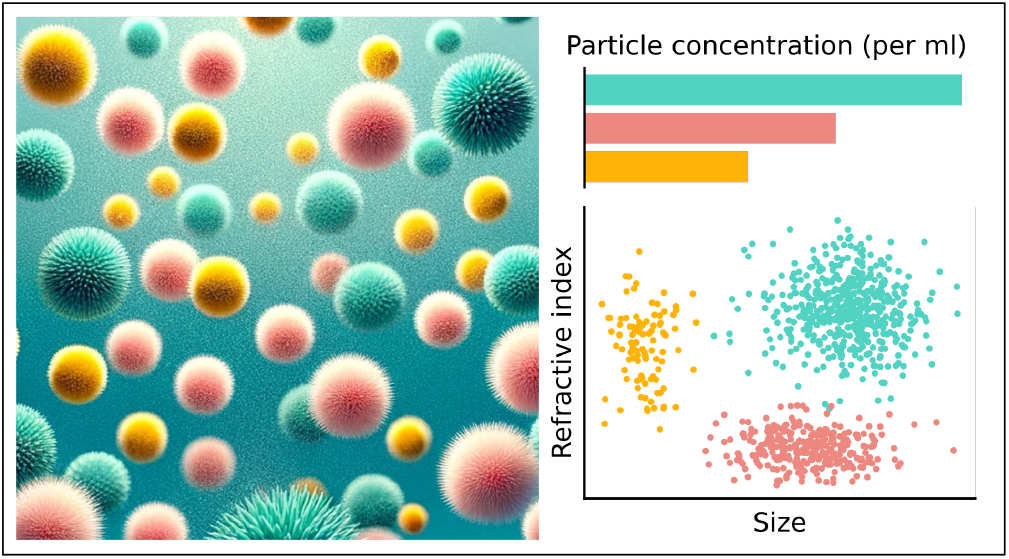

